# Development of a Synthetic Biosensor for Chemical Exchange MRI Utilizing *In Silico* Optimized Peptides

**DOI:** 10.1101/2023.03.08.531737

**Authors:** Adam J. Fillion, Alexander R. Bricco, Harvey D. Lee, David Korenchan, Christian T. Farrar, Assaf A. Gilad

## Abstract

Chemical Exchange Saturation Transfer (CEST) magnetic resonance imaging (MRI) has been identified as a novel alternative to classical diagnostic imaging. Over the last several decades, many studies have been conducted to determine possible CEST agents, such as endogenously expressed compounds or proteins, that can be utilized to produce contrast with minimally invasive procedures and reduced or non-existent levels of toxicity. In recent years there has been an increased interest in the generation of genetically engineered CEST contrast agents, typically based on existing proteins with CEST contrast or modified to produce CEST contrast. We have developed an *in-silico* method for the evolution of peptide sequences to optimize CEST contrast and showed that these peptides could be combined to create *de novo* biosensors for CEST MRI. A single protein, superCESTide 2.0, was designed to be 198 amino acids. SuperCESTide 2.0 was expressed in *E. coli* and purified with size-exclusion chromatography. The magnetic transfer ratio asymmetry (MTR_asym_) generated by superCESTide 2.0 was comparable to levels seen in previous CEST reporters, such as protamine sulfate (salmon protamine, SP), Poly-L-Lysine (PLL), and human protamine (hPRM1). This data shows that novel peptides with sequences optimized *in silico* for CEST contrast that utilizes a more comprehensive range of amino acids can still produce contrast when assembled into protein units expressed in complex living environments.

## 2 Introduction

The development of genetically engineered reporters to enhance various imaging modalities has led to the elucidation of many biological mechanisms. With a wide range of amino acids, the ability to engineer synthetic proteins with different functionality is near endless, with even more possibilities generated by the creation of non-canonical amino acids. One of the imaging modalities that genetically engineered reporters have enhanced is Magnetic Resonance Imaging (MRI). Several genetically encoded sensors have been developed based on different MRI contrast mechanisms^1,2^, such as transverse relaxation^3–6^, longitudinal relaxation^7–9^, and chemical exchange saturation transfer (CEST)^10–18^.

CEST MRI has recently taken center stage due to the rapid development of CEST methodologies^19,20^. A significant advantage of CEST MRI is that it allows for the detection of contrast based on natural metabolites such as amino acids^21^, peptides^22–24^, and sugars^26,26^ along with sensitivity to pH^27–29^, making it suitable for clinical imaging. CEST MRI is also a powerful tool for noninvasive molecular imaging of paramagnetic organic compounds^31,32^, but primarily for diamagnetic metabolites^33^ and peptides^34^. In our lab, research has been conducted to create an *in-silico* method for the directed evolution of short peptide sequences termed Protein Optimization Engineering Tool (POET). While developing POET, the algorithm was trained to produce a library of synthetic peptides to generate contrast utilizing CEST MRI. There was particular interest in CEST due to the novelty of the mechanisms involved and the versatility of its application which allowed for an extensive range of peptides to be tested.

POET outperformed expectations and produced peptides up to 4 times stronger than the original PLL test data with a more diverse amino acid range. Still, concerns existed around the *in vivo* applications of these peptides. When these completely synthetic peptides were resuspended in controlled, buffered environments, the contrast produced was significant. However, if these peptides were expressed as small genetic units in a living environment, would the CEST properties of the peptides be altered, eliminating any of the advantages gained through *in-silico* evolution? The answer to such a question could be answered by looking at the peptides through the lens of proteomics and understanding them structurally.

The field of proteomics has grown tremendously over the last couple of decades and has provided a strong foundation for the engineering of synthetic peptides and proteins. De *novo* amino acid synthesis has simplified the screening of large peptide libraries for desired structures, functions, and assemblies^35^. Still, this synthesis method lacks the *in vitro* and *in vivo* considerations involved in the design of therapeutic biomolecules. Studies have shown that careful considerations need to be made during therapeutic protein design due to degradation pathways and drastic changes in environmental conditions that are present *in vivo*^36^. Under these physiological conditions, the CEST contrast of these 12-mer peptides could be degraded, and the pH could alter the interactions of the individual amino acids. It’s widely understood that the generation of secondary and tertiary structures can lead to greater thermal protein stability^37^. Still, these structures are only accessible to more extensive amino acid sequences due to the increased complexity of intramolecular interactions^38^. These characteristics common to large protein sequences presented a seemingly simple solution to the problem faced with the small CEST peptides. By combining the peptides from the previously generated library into larger proteins, secondary and tertiary structures could form, making them more robust and apt to survive in a complex living environment.

Through the following study, we sought to combine these peptides into a single genetically encoded unit. We examined if CEST contrast could still be generated when combined as a larger unit and if the complex environments of living organisms were amenable to the introduction of CEST proteins. Our findings show that a gene encoding a synthetic polypeptide, termed superCESTide 2.0, could be expressed and purified from *E. coli*. SuperCESTides can generate clear differences in CEST contrast compared to other proteins. Moreover, superCESTide 2.0 has a more diverse amino acid composition that reduces the burden on the cellular metabolism, thus potentially allowing the production of higher reporter concentrations and reducing the risk of mutagenesis^39^.

## 3 Experimental Methods

### 3.1 Gene Synthesis and Cloning

Using the previously generated peptide library targeted for 3.6-ppm contrast from the Protein Optimization Engineering Tool^40^, an amino acid sequence was generated by connecting the top-performing peptides end to end to create a 198 (superCESTide 2.0) amino acid long protein. The amino acid sequence was then optimized for DNA expression in *E. coli* using the Azenta Life Sciences’ Codon Optimization tool on Genewiz. The sequence was cloned using TOPO cloning, first in silico into the pET101 expression plasmid using SnapGene’s directional TOPO cloning simulation. DNA sequences were then purchased as G-Blocks from Integrated DNA technologies. TOPO cloning was performed using Champion pET101 Directional TOPO expression Kit (ThermoFisher, K10101). pET101 recombinant plasmids were transformed into One Shot TOP10 Chemically Competent *E. coli* (ThermoFisher, C404010). Bacterial colonies were grown on μg/mL ampicillin-rich agar plates overnight at 37 °C. Colony PCR was performed using Quick-Load *Taq* 2X master mix (New England Biolabs, M0271L). Successful colonies were grown in 100 μg/mL ampicillin-rich lysogeny broth (LB) overnight at 37 °C. DNA was extracted using the QIAprep Spin Miniprep Kit (Qiagen, 27106X4) and then sent to Azenta Life Sciences for DNA sequencing. Sequencing data were then aligned with the *in silico* pET101 recombinant plasmids to verify successful cloning.

### 3.2 BL21 *E. coli* Expression

Successfully cloned recombinant pET101 plasmids were transformed into One Shot BL21(DE3) Chemically Competent *E. coli* (ThermoFisher, C600003) and plated on 100 μg/mL ampicillin-rich agar plates overnight at 37 °C. A screen of different growth conditions was performed to determine optimal culture conditions. IPTG induction was performed by inoculating 2 mL ampicillin-rich LB media with a single recombinant BL21 *E. coli* colony and growing the culture at 37 °C until an OD_600_ of 0.6 was reached. Bacteria were then induced with 0.5 mM IPTG and grown at 37 °C, 30 °C, and 22 °C for 18 hrs. Cultures of recombinant BL21 *E. coli* were also grown in 2 mL Overnight Express Instant TB media (Sigma Aldrich, 71491-5) with added 100 μg/mL ampicillin; cultures were grown at 37 °C, 30 °C, and 22 °C for 18 hrs. A single culture of ampicillin-rich LB media was inoculated with recombinant *E. coli* but left uninduced to act as a negative control for expression. Protein was extracted using BugBuster Master Mix (Sigma Aldrich, 71456), mixed 1:1 with 2x laemmli sample buffer (Sigma Aldrich, S3401), incubated at 95 °C, and then run on an Any kD Mini-PROTEAN TGX Stain-Free Protein gel (Biorad, 4568124). SDS-PAGEs were captured on the Biorad Chemidoc Imaging System. Immunoblot was performed by transferring protein to a PVDF membrane using Trans-Blot Turbo RTA Transfer Kit (Biorad, 1704272) and the Biorad Transblot Turbo System. PVDF membrane was blocked with 5% milk and stained with monoclonal anti-polyHistidine primary antibody (Sigma Aldrich, H1029) and polyclonal anti-mouse secondary antibody, and imaged with Clarity Western ECL substrate (Biorad, 1705061) and the Biorad Chemidoc Imaging system.

### 3.3 Size Exclusion Chromatography, Fraction Identification

To perform Size Exclusion Chromatography, the Äkta Start chromatogram system was utilized from cytiva paired with the Hiprep 16/60 Sephacryl S-200 HR (Cytiva, 17116601) to perform size exclusion chromatography (SEC). To calibrate the instrument and determine a procedure for SEC, gel filtration standards (Biorad, 1511901) were used to create a calibration curve with the following chromatogram parameters: Flow rate of 0.5 mL/min, max pressure of 0.15 MPa, room temperature, 100% phosphate buffer saline mobile phase, elution volume equal to one column volume (120 mL). To identify regions of the chromatogram where the synthetic protein was eluting, thirty 1 mL fractions were collected around the estimated molecular weight. Fractions were precipitated by adding −20 °C acetone at a volume 4 times greater than the original fraction. Samples were then vortexed briefly and incubated at −20 °C for 1 hour. Protein was pelleted by centrifuging the sample at 15,000 x g for 10 min. The acetone was poured off, and the pellet was allowed to dry for 15 minutes to remove excess acetone. The semi-dry pellet was then resuspended in PBS at a volume 10x smaller than the original fraction (100 μL). Dot blots were performed using a PVDF membrane that was first wetted with 100% ethanol until the membrane was transparent. The membrane was then incubated in a transfer buffer from the trans-blot turbo RTA transfer kit (Biorad, 1704272) for 5 minutes. While incubating, tween 20 was added to the transfer buffer to create a 0.5% (v/v) solution (TBS-T). The membrane was transferred to a dry dish and semi-dried before pipetting 2 μL of each sample onto the blot. Dots were allowed to dry, and then the membrane was blocked with 5% milk in TBS-T for 1 hour at room temperature. A primary antibody solution was made with monoclonal anti-polyHistidine antibodies (Sigma Aldrich, H1029) diluted 1:1000 in 5% milk TBS-T. The blocking buffer was removed from the dish, and the primary antibody was poured over the membrane. The membrane was incubated for 1 hour with constant motion at room temperature or overnight with constant motion at 4 °C. Once the incubation was finished, the membrane was washed with TBS-T three times, for 5 min each. During washes, a 1:10000 secondary antibody solution was made by diluting polyclonal anti-mouse antibodies (Sigma Aldrich, A4416) in TBS-T. After washing the membrane, the secondary antibody solution was added and incubated at room temperature with constant motion for 1 hour. Once the incubation was complete, the membrane was washed twice with TBS-T for 5 minutes and washed a third time with TBS for 5 min. The membrane was imaged with clarity western ECL substrates (Biorad, 1705061) and the Biorad Chemidoc imaging system.

### 3.4 Size Exclusion Chromatography, Sample Collection

Expression was carried out to collect soluble, non-denatured CEST proteins as described in section 3.2. Size exclusion chromatography was performed with the same procedure created for fraction identification described in section 3.3. Fraction identification showed that superCESTide 2.0 eluted at 52 mL to 60 mL and human protamine eluted at 44 mL to 55 mL from the Hiprep 16/60 Sephacryl S-200 HR column. Repeated separations were performed, and the same region of fractions was collected and concentrated using Amicon Ultra – 15 centrifugal filters, 10 kD (Sigma, UFC9010). Samples were resuspended in phosphate-buffered saline and stored at 4 °C until all samples were ready for MRI analysis. The protein concentration of the samples was determined using Pierce Coomassie Plus (Bradford) Assay kit (Thermofisher, 1856210). Sample proteins were normalized to the lowest concentration sample using phosphate-buffered saline.

### 3.5 CEST MRI Parameters

CEST scans were acquired using a horizontal bore 7T Bruker preclinical MRI with Paravision v3.1. The samples were placed within a custom-designed imaging phantom produced by 3D printing specifically for this task. The phantom allows simultaneous imaging of 12 samples at 200 μL per well. A single set of scans was comprised of two different pulse sequences. The first scan is a WASSR scan used to determine the exact frequency of water in the sample so that it may be adjusted accordingly^41^. The second scan is a CEST scan made from a modified RARE sequence, with a RARE factor of 25 and a TR of 12,000 ms. Saturation pulses were applied as a block pulse for 4,000 ms, and a saturation power of 3 μT covering saturation frequencies from −7 to 7 ppm offset from water in steps of 0.1 ppm. During the scans, the samples were heated by controlled air flow connected to the sample platform and held at a constant temperature of 37 °C. Each set of samples was scanned three to five times to perform statistical analyses. Data processing was performed with a custom MATLAB script^42^.

## 4 Results and Discussion

### 4.1 Design of synthetic genes - superCESTides

The high percentage of lysine and arginine residues in genetically encoded CEST-based reporters^11,13,15,18^ burden cellular metabolism. Consequently, this burden on metabolism reduces the reporter protein’s cellular concentration, leading to a decrease in overall contrast. To remedy this issue, we developed a machine learning algorithm that evolved peptides to optimize their CEST contrast while diversifying the amino acid residues in their sequences^40,43^. Having generated a wide range of 12 amino acid long peptides, questions about their expression in cells still existed. While these *de novo* peptides produced relatively high CEST contrast when suspended in solution, it was unknown if they would still create detectable contrast once expressed as a single genetic unit in a complex organism. To address these concerns, it was hypothesized that these peptides could be assembled into larger proteins to increase stability by the resulting secondary and tertiary structures while enhancing the contrast being produced. This assembly of peptides would simultaneously address the burden on cellular metabolism and stability by diversifying the amino acids in the sequence. To assemble this synthetic CEST reporter, the strongest 3.6 ppm contrast-producing peptides were joined end to end (Table S1). These sequences were analyzed for repeated sequences of amino acids that could lead to hairpin formation. An artificial protein, encoded by a single independent gene, was termed superCESTide 2.0, a 198 amino acid reporter (Fig. S1). Figure 1 shows the resulting distribution of amino acids in the newly created superCESTide 2.0 compared to previous CEST reporters and the predicted structure of this synthetic protein.

**Figure S1.**
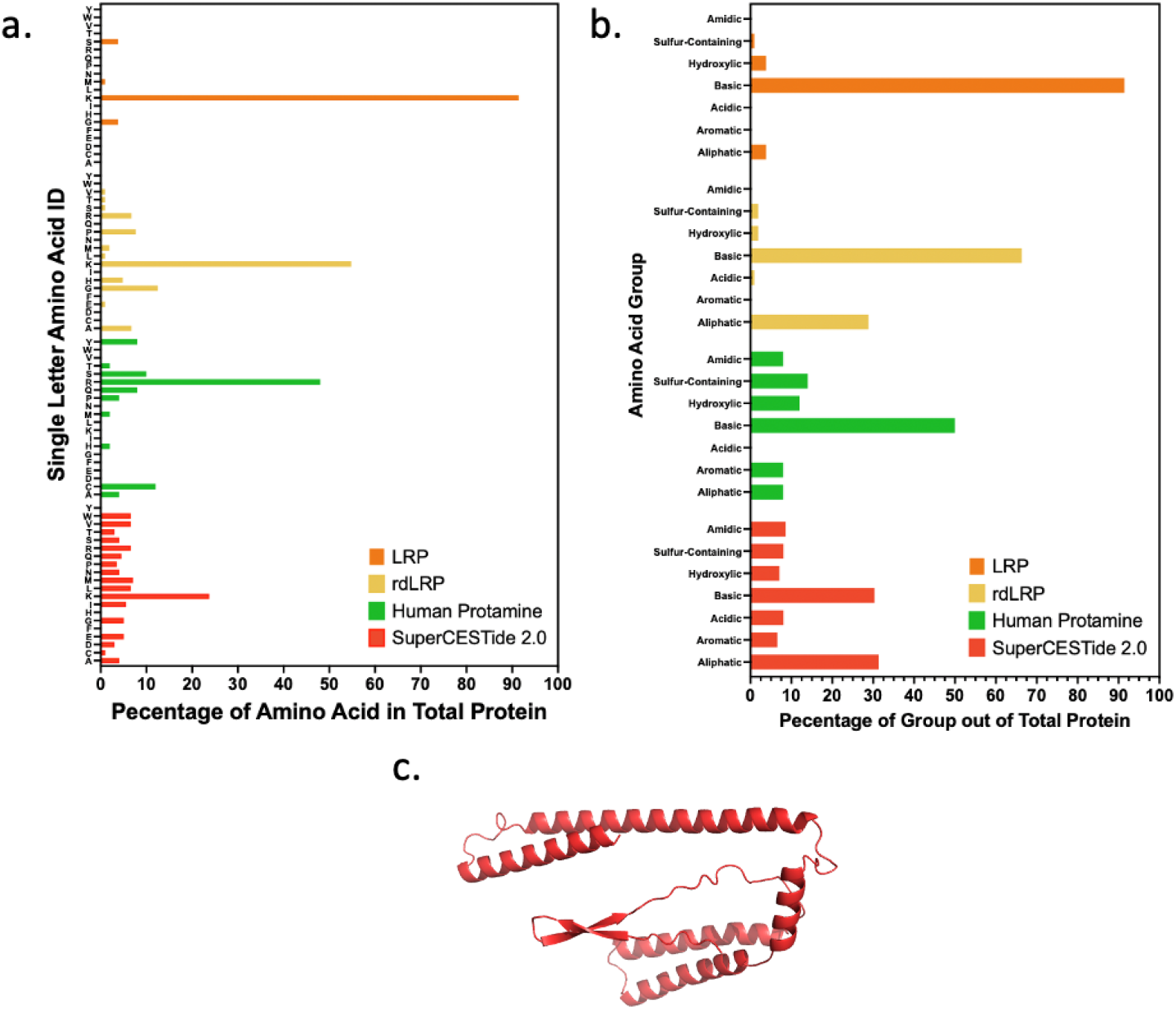
Percentages of Amino Acid Composition in Previous CEST Reports Compared to New Synthetic CEST Reporters. **(a)** The bar graph shows each amino acid’s portion of the total protein as a percentage. A specific color displays each reporter: orange representing LRP^8^, yellow representing rdLRP^10,47^, green representing hPRM1^15^, and red representing superCESTide 2.0. **(b)** The bar graph shows each amino acid group’s portion of the total protein as a percentage. Amino acids were grouped based on the properties of their associated side chains. Each reporter is displayed by a specific color, with orange representing LRP, yellow representing rdLRP, green representing hPRM1, and red representing superCESTide 2.0. **(c)** Predicted structure of superCESTide 2.0 generated by FoldX and displayed in PyMOL.

As seen in figure 1, the diversity of amino acids in early reporters was minimal, with a significant focus on basic residues, specifically lysine and arginine. LRP, one of the old synthetic reporters we studied, was dominated by the presence of lysine, making up more than 90% of its sequence, making it incredibly burdensome for synthesis and could limit expression in living hosts. When looking at the superCESTide 2.0, the distribution of amino acids is much greater than in previous reporters, limiting the strain on host cells and allowing for a broader range of applications. However significant these advantages may have seemed, having only generated these structures *in silico* didn’t illustrate the properties that superCESTide 2.0 provided as a reporter, and the next step would be to express these genes in *E. coli*, such that the artificial protein could be purified and screened for CEST contrast in a simple and controlled environment.

### 4.2 Crude Purification of superCESTide 2.0 Using Amicon Centrifugation Filters

To analyze the CEST contrast generated by superCESTide 2.0, it would first need to be expressed in *E. coli* and the protein removed by lysis so it could be scanned by MRI. Whole-cell lysates contain a myriad of proteins and lipids, and trace detergents used for lysis, all of which can contribute to the contrast observed in CEST MRI. To determine if the contrast produced was due to the presence of superCESTide, a purification method would need to be utilized to remove any contributing particles, such that the subtle differences could be attributed solely to the recombinant proteins present in each sample.

Of the purification methods considered, the most logical step was the utilization of cobalt-resin-based affinity chromatography to target the 6HIS-tag attached to the c-terminus of each superCESTide. However, as seen in Figure S2, we had little retention of target protein, with no significant contribution to contrast following affinity purification. Thus, we searched for alternative methods and conceptualized a technique for utilizing amicon centrifugation filters as a means of purification. The marketed use of amicon centrifugation filters is to retain a target protein with a molecular weight greater than the molecular weight cut-off of the filter being used while allowing smaller contaminating molecules to pass through with the excess buffer or solvent. Theoretically, a filter with a molecular weight cut-off of 50 kDa, almost twice the size of our target protein, superCESTide 2.0, would allow our target to pass through into the filtrate. The filtrate could then be concentrated by a filter with a molecular weight cut-off of 10 kDa (Figure 2a). Before attempting this with our synthetic protein, the process was initially tested and optimized with a protein of similar molecular weight, super folding Green Fluorescent Protein (sfGFP), which would simplify the visualization of protein passing through the filter by utilizing its fluorescence. Once a protocol was developed for separating sfGFP, it was reapplied to our synthetic reporters. The SDS-PAGE in Figure 2b shows the results of the crude purification of superCESTide 2.0, with each lane displaying the proteins present at the various stages of purification. After filtration and concentration, the fraction containing superCESTide 2.0 showed a clear band on the SDS-PAGE (Figure 2b, lane 4) at a molecular weight previously determined by a western blot of soluble superCESTide 2.0 (Figure S2a, MW ~25 kDa).

**Figure 2.**
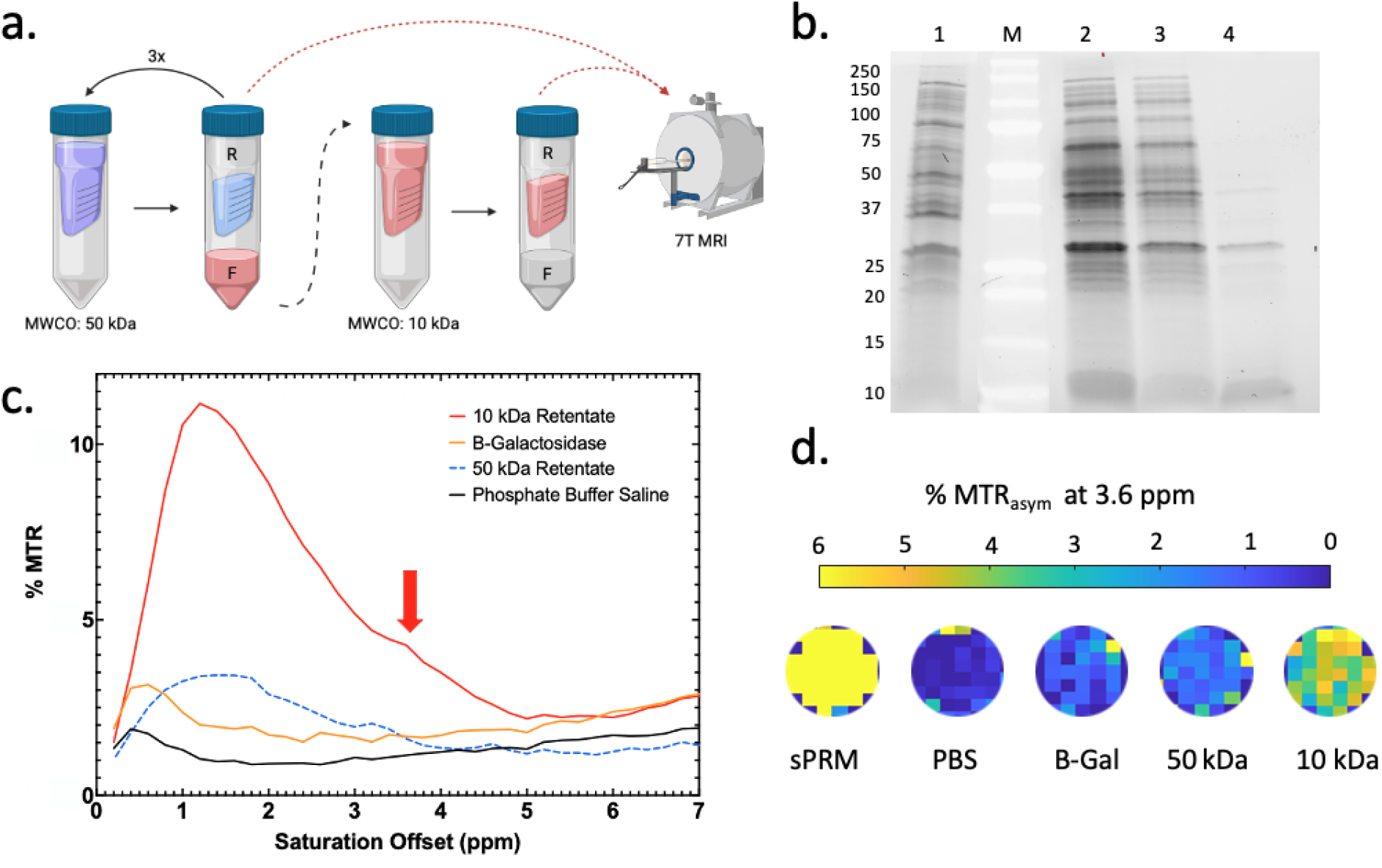
Crude purification of superCESTide 2.0 with amicon centrifugal filters. **(a)** Diagram showing the steps taken to purify the protein, the purple solution represents the bacteria lysate containing superCESTide 2.0, the blue solution represents the proteins retained by the 50 kDa filter, and the red solvent represents the flow through containing the crudely purified superCESTide 2.0. R and F indicate the retentate and filtrate, respectively, (b) SDS-PAGE of samples obtained while purifying superCESTide 2.0 using amicon centrifugal filters. Lane 1: Insoluble fraction of superCESTide 2.0 diluted 1:10, Lane M: Molecular Weight Markers, Lane 2: Unfiltered soluble fraction of superCESTide 2.0, Lane 3: 50 kDa retentate of soluble superCESTide 2.0 diluted 1:10, Lane 4: 10 kDa retentate of soluble superCESTide 2.0 diluted 1:10. **(c)** MT_Rasym_ plot of superCESTide 2.0 purified using amicon centrifugal filters. 50k retentate of superCESTide 2.0 was all normalized to the protein concentration of the 10k retentate (1.39 mg/mL). The solid red line represents the retained protein from the 10 kDa amicon centrifugation filter, the solid orange line represents purified β-gal, and the dashed blue line represents all the protein retained by the 50 kDa filter. The red arrow points to the amide peak at 3.6 ppm. (d) ROIs of each well on phantom from MTR_aysm_ map generated by MRI, the max MT_Rasym_ at 3.6 ppm is 6% with yellow being the most intense signal.

Next, the 50 kDa and 10 kDa retentates were imaged using MRI. To ensure the change in contrast in the superCESTide 2.0 sample was due to the presence of the CEST reporter, a protein known not to produce contrast at 3.6 ppm, in this case, β-galactosidase (β-gal) was also transformed into bacteria and purified so it could be ensured that the introduction of a recombinant plasmid with a foreign protein wasn’t enough to alter the CEST contrast produced.

As seen in in figure 2, the 10 kDa retentate showed an MTR_asym_ curve with a peak forming at 3.6 ppm. This was the first indication that these fractions may contain soluble superCESTide 2.0 and that the contrast generated may be a direct result of the presence of SuperCESTide. Figure 2c further supported this observation with the lack of contrast generated by the proteins in the 50 kDa retentate, especially around the amine and the amide resonance frequency. Notably, the small peak at 3.6 ppm is characteristic of fast-exchanging amide protons previously observed in the peptides optimized through the protein engineering tool. This is usually the hallmark of CEST-based reporters and was the first indication that the assembled peptides could produce contrast above background levels when expressed as a larger protein in a complex living organism. Looking back at the SDS-PAGE in Figure 2b, it was observed that a large portion of our target protein was retained in the 50 kDa filter, and there was concern regarding the scalability of the amicon purification. Other methods were investigated for separation at comparable levels of specificity to the amicon filters.

### 4.3 Size Exclusion Chromatography for Purification

In principle, the method developed by utilizing amicon centrifugation filters with differing molecular weight cutoffs was similar to size exclusion chromatography (SEC), where molecules could be separated based on their interaction with a stationary matrix relative to their molecular weight. An SEC column was obtained for the chromatographic system, Aktä Start, to scale up the purification and decrease the amount of synthetic reporter lost during purification. The column was calibrated with gel filtration standards and then used to separate our synthetic protein. Initial chromatographic separations based on calibration curves alone were unsuccessful (Figure S3), so a different method for protein identification needed to be developed. 1 mL fractions were collected around the theoretical molecular weight of superCESTide 2.0. The protein in each fraction was concentrated with acetone precipitation and resuspended in a volume 10-fold smaller than the original fraction. Each fraction was then screened for the presence of superCESTide 2.0 by performing a dot blot with anti-6HIS antibodies, and the results can be seen in figure 3. The proteins collected in the fractions ranged from 14 kDa to 130 kDa (Figure 3a), encompassing our previously determined molecular weight (Figure S2a). All thirty fractions were precipitated and screened with a dot blot. In figure 3b, the dots from the dot blot were cropped and placed in the order in which they were collected. The region that displayed binding after incubation with anti-6HIS antibodies was highlighted in grey in figure 3. With fractions containing superCESTide 2.0 identified, a second separation was performed to collect concentrated protein with amicon filters instead of utilizing acetone precipitation to ensure that the denaturation didn’t alter the properties of the synthetic CEST reporter.

**Figure 3.**
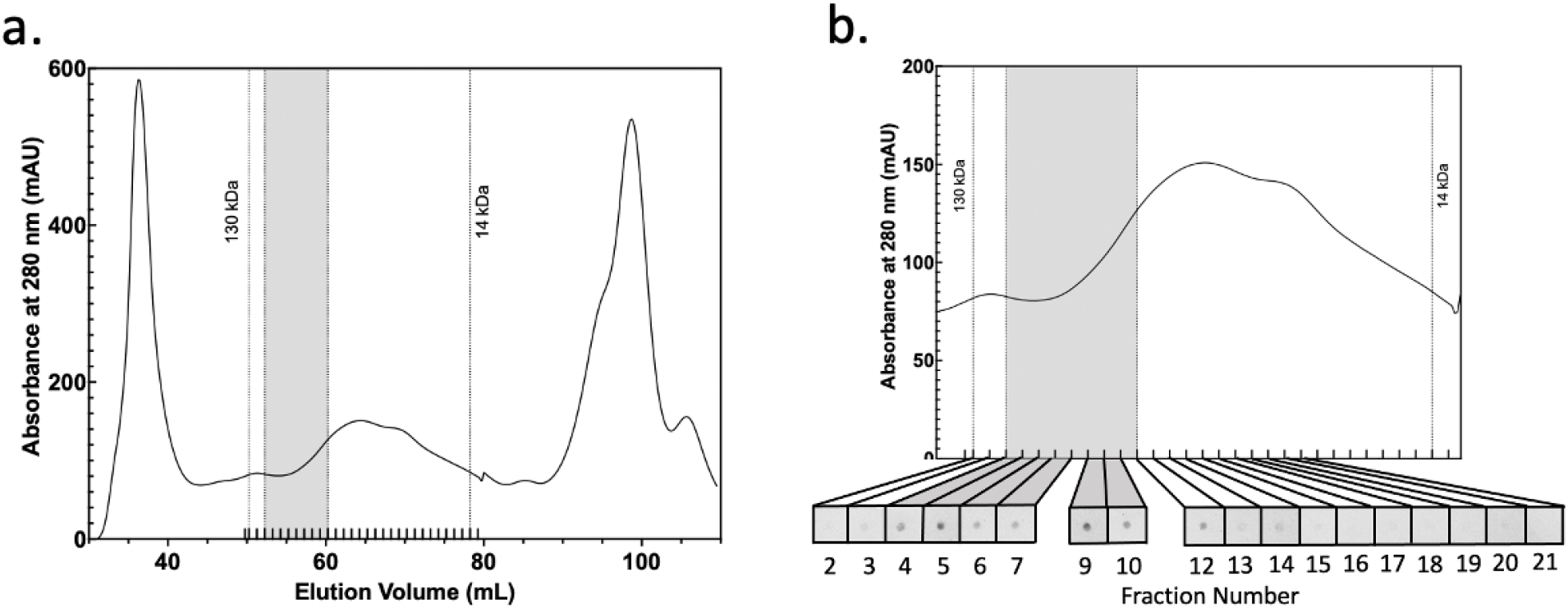
Purification and Identification of superCESTide 2.0 with size exclusion chromatography. **(a)** Chromatogram of superCESTide 2.0 collected for concentrated natured superCESTide 2.0 protein. The grey region shows the fractions that were collected for concentration. The dashed lines encompass the area that was collected with the fractionator, **(b)** The expanded area of the chromatogram collected by the fractionator. A dot blot of superCESTide 2.0 can be seen below the chromatogram with dots in order of their fraction number.

With SEC, the separation and identification of SuperCESTide 2.0 were successful, but it was understood that SEC separation only removed proteins outside of a particular range of molecular weights. This meant that the separated superCESTide 2.0 sample still contained endogenous *E. coli* proteins similar in size to superCESTide 2.0, which could contribute to the overall contrast. To address concerns regarding the non-specific nature of SEC and the contribution of contaminating proteins, a third separation was performed on a culture expressing a plasmid containing bovine pancreas trypsin inhibitor (BPTI), a known protein included in protease inhibitor cocktail used during bacterial cell lysis. These lysates were separated with the same protocol used for superCESTide 2.0, and the same region of the eluate was collected. These resulting fractions contained proteins that eluted at the same volume as superCESTide 2.0 and would serve as a blank. The resulting contrast could then be attributed to the presence of the CEST reporter protein. The superCESTide 2.0 and blank samples obtained from SEC were then scanned by MRI. Figure 4 shows the MTR_asym_ of superCESTide 2.0 SEC purification compared to the blank sample containing BPTI annotated as superCESTide 2.0 blank.

**Figure 4.**
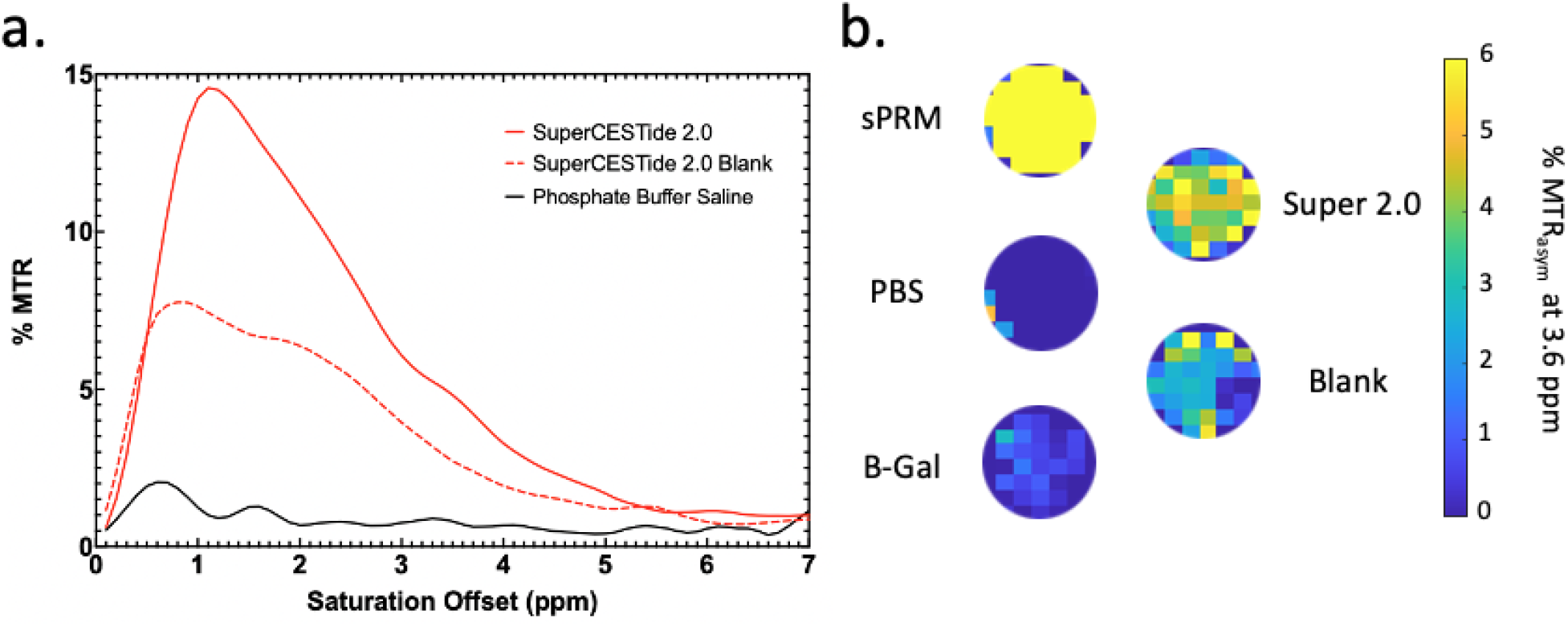
MTR_aS_ym plot of SuperCESTide 2.0 compared to the superCESTide 2.0 blank. **(a)** The blue curve represents salmon protamine, a control for contrast at 3.6 ppm. The solid red line represents superCESTide 2.0 following purification with SEC and concentration with amicon filters, and the dashed red line represents the blank fractions where superCESTide 2.0 elutes. The red arrow points to the amide peak at 3.6 ppm. (b) ROIs of each well on phantom from MTR_aysm_ map generated by MRI, the max MTR_asym_ at 3.6 ppm is 6% with yellow being the most intense signal.

As seen in figure 4, the superCESTide 2.0 sample generated much higher CEST contrast with distinct peaks at the typical resonance frequency of the exchangeable amide, and amine protons (Figure 4a) compared to the lack of contrast produced by the blank. As seen from previous purification methods (Figure 1a, Figure 2c), the same distinct peak formed at 3.6 ppm following SEC. These findings further support the hypothesis that a recombinant protein composed of multiple peptides optimized for generating contrast at 3.6 ppm can generate measurable CEST contrast exceeding the CEST contrast produced by *E. coli’s* endogenous cellular components.

### 4.4 Comparison of SuperCESTide 2.0 to Prior CEST Reporters

Through this study, we attempted to understand the limits involved in synthetic protein amino acid composition and how it played a role in designing novel CEST biosensors. Our results indicate that *in silico* peptide optimization tools can generate novel proteins from peptides utilizing a wide range of amino acids that still produce CEST contrast while limiting the effects of single codon usage (Figure S4), which can lead to strain on metabolic pathways^39^. Through the successful expression of superCESTide proteins in *E. coli* and further identification and measurement of contrast, we demonstrated the amenability of these natural living environments to the presence of synthetic CEST agents. The ability of these superCESTides to produce CEST contrast following transformation in *E. coli* further illustrates the resilience of CEST reporters in a complex environment through the possible presence of secondary and tertiary structures in the protein and interactions with other endogenously expressed proteins. To demonstrate the efficacy of this new reporter, we wanted to compare this synthetic reporter to a human-based protein reporter previously identified by our group, human protamine (hPRM1)^15^. Both hPRM1 and superCESTide 2.0 were expressed and separated using size exclusion chromatography, and the concentration of the samples was normalized to the lowest sample before MRI scans. Figure 5 shows the results following three consecutive SEC separations of hPRM1 and superCESTide 2.0, yielding a final concentration of 6.6 mg/mL protein. Figure 5 also includes protamine sulfate, a CEST contrast agent, at a 2.5 mg/mL concentration. The inclusion of protamine sulfate was due to its characteristic MTR_asytγi/_ which illustrates the ideal amide contribution sought in a CEST reporter. Protamine sulfate, being commercially purified, contains no contaminating proteins and therefore was scanned at a lower concentration than the SEC samples to compensate for the contribution of contaminants to the SEC sample’s protein concentrations. Due to the uncertainty of the exact concentration of superCESTide 2.0 and hPRM1 in the SEC samples, any attempt to directly compare them to commercially purified protamine sulfate would be inaccurate. Still, it serves to represent the ideal MTR_asym_ for CEST reporters with large amide proton contribution.

**Figure 5.**
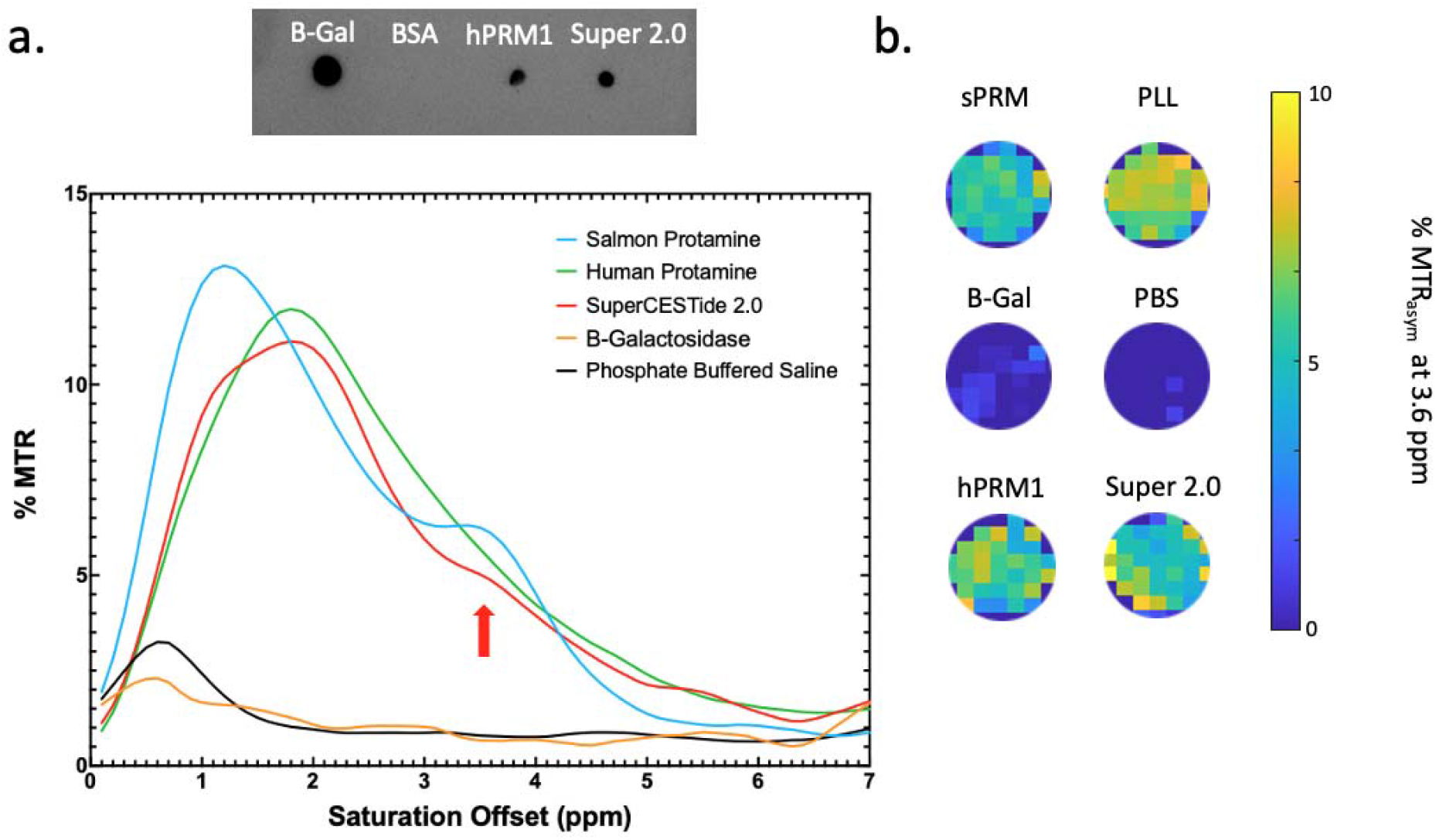
MTR_aS_ym Plot comparing hPRM1 and SuperCESTide 2.0 following SEC. **(a)** Dot blot of hPRM1 and superCESTide 2.0 samples following SEC with β-galactosidase (β-gal) as a positive control for 6HIS+ protein, bovine serum albumin (BSA) serving as a negative control for 6HIS-protein. On the MTR_asym_ plot, Salmon Protamine (sPRM) is represented by the blue line, the green line represents hPRM1, the orange line represents β-gal, and superCESTide 2.0 is represented by the red line. The legend is ordered by contrast generated at 3.6 ppm, with most contrast generated listed at the top. The red arrow points to the amide peak at 3.6 ppm. **(b)** ROIs of each well on phantom from MTR_aysm_ map generated by MRI, the max MTR_asym_ is 10% with yellow being the most intense signal.

As seen in Figure 5, all the reporters produced comparable levels of contrast, and superCESTide 2.0 produced a more distinctive amide peak at 3.6 ppm than hPRM1. This further supports the ability to create CEST contrast in a synthetic protein and shows that these synthetic proteins are as effective as previously identified reporters. Despite similar levels of contrast, the distinct advantage of superCESTide 2.0 over hPRM1 and protamine sulfate is the variety of amino acids present in the sequence. If these biological reporters are to be used for imaging in tumor microenvironments, the constraints of these environments need to be considered. Previous studies have shown that tumors rapidly mutate to overcome the limitations of cellular metabolism, and one such limitation that tumors have overcome is the bioavailability of arginine^39^. Having a biological reporter, such as hPRM1, heavily reliant on a single amino acid codon could be limiting in extreme cellular environments and reduce the effectiveness of the imaging reporter. Such concerns are reduced in the synthetic superCESTide 2.0 due to the variety of amino acids present, allowing for a more extensive range of imaging applications. Even though the generation of contrast from a synthetic protein was successful, the investigation of superCESTide 2.0 as a biosynthetic reporter is still ongoing.

The difficulties experienced during the purification of superCESTide 2.0 pose further avenues of study. Due to the utterly synthetic nature of these proteins, more information is needed to understand their structures, and few tools exist that can predict these structures with high confidence levels. The increase in amino acid residues is believed to increase the likelihood of secondary and tertiary structures through intramolecular amino acid side chain interactions. Still, the complexity of signaling involved in post-translational modifications may limit such structures from forming. Future studies into the characterization of these superCESTides will need to be performed to understand their presence in the cytoplasm. Their overall stability and elucidation in structure could aid in the development of more effective methods for purification and downstream processing.

As has been the interest with previous CEST reporters, the goal for these superCESTides is to eventually utilize them for clinical diagnostics. The lack of knowledge about synthetic proteins leaves many avenues for investigation. One major hurdle for clinical trials will be the characterization of superCESTide immunogenicity. It is well understood and characterized that the presence of foreign proteins in the body will eventually illicit an immune response, either from antigen-presenting cell (APC) MHC class II molecules following introduction into the body^44^ or through the presentation in MHC class I molecules following transduction or transfection of nucleated cells^45^. Prior studies of CEST reporters conducted by our lab and others indicate that there appear to be limited effects *in vivo*. However, a long-term study would need to be undertaken to understand the implications of the immune system’s reactivity to these reporter proteins.

Nevertheless, the finding from this study strongly supports the notion that immunogenetic epitopes could be replaced with other peptides while retaining CEST characteristics. It might be feasible to expand the POET to include a module that screens for immunogenic epitopes using known algorithms and exclude them, much like we excluded hydrophobic peptides to humanize the CEST reporters. Though, in general, the CEST-based reporters are intended to be expressed in the cytoplasm and, thus, slow the initial exposure to the host immune system, understanding methods for immune compliance or avoidance may be pivotal in bolstering the efficacy of these CEST peptides as MRI reporters. Future clinical trials could include a series of CEST peptides, each producing novel antigens, to limit the immune response of chronic MRI patients. In addition, investigating and elucidating the limited immune response to human placental cells^46^ could provide unique insights into future diagnostic applications of these CEST reporters. It is also possible that immune response may not be a limitation for some pre-clinical studies, such as in cases where immune responses might be beneficial, like oncolytic virotherapy.

Regardless of concerns around the future applications of these synthetic proteins, a significant step has been taken in developing *de novo* synthetic proteins for diagnostics, and the limitations of previous CEST reporters have been overcome. Through further optimization of *in silico* peptides and the elucidation of structural characteristics, future generations of superCESTides could see resolution in the issues currently faced with purification and immunogenicity and further enhance their diagnostic capabilities by further improving CEST contrast signal.

## 5 Conclusion

This study demonstrated that superCESTide, a synthetic gene assembled from monomeric peptides optimized by a genetic programming algorithm, can produce marked CEST contrast when expressed in bacteria and perform at levels comparable to previously identified reporters. Due to its diverse amino acid composition compared to previous generations of CEST-based reporter genes^10,14,15,18^, superCESTide can improve DNA stability and reduces the burden on cellular protein translation and metabolism. Moreover, the interchangeable monomers of the sequence can open the possibility of enhancing the immunocompatibility of the superCESTides by replacing potentially immunogenic epitopes.

## List of Abbreviations

CEST: Chemical Exchange Saturation Transfer
β-gal: β-galactosidase
MTR_asym_: Magnetic Transfer Ration Asymmetry
SP: Salmon Protamine, Protamine Sulfate
PLL: Poly-L-Lysine
hPRM1: Human Protamine 1
LRP: Lysine Rich Protein
TBS-T: Transfer buffer solution + 0.5% Tween 20
POET: Protein Optimization Engineering Tool
SEC-Size: Exclusion Chromatography
sfGFP: Super Folding Green Fluorescent Protein
BPTI: Bovine Pancreas Trypsin Inhibitor
APC: Antigen Presenting Cells
MHC: Major Histocompatibility Complex

## Acknowledgments

The author would like to acknowledge financial support from the NIH/NINDS: R01-NS098231; R01-NS104306 NIH/NIBIB: R01-EB031008; R01-EB030565; R01-EB031936; P41-EB024495 and NSF 2027113.

