## Supplemental Information for "Development of a Synthetic Biosensor for Chemical Exchange MRI Utilizing *In Silico* Optimized Peptides"

**6 – Supporting Information**

**6.1 – Synthetic Reporter Sequences**

SuperCESTide 2.0 DNA Sequence:

ATGCTGTGGAGCGATATTAAAATGAAACTGAAAAAAACCAAAATGGGCAAACTGATTGGCATTCCGGTGCTGAAAATGAAAGTGGCGGCGGCGATGGCGCCGAAACAAGTGAAGATGTGGGACTGGGAACAAAAAAAGAAGTGGATTCGACTACCGAAACGCGTGCAAGGCAACGTGGAAAAAGTGACCCGCATGACCATTCAAGTGAAAGGCAGCAAAGCGAAATGCAAAGTGCAGAGCGCGAACGTGTGCAAACCGGTGAACCGCCTGGGCAAAATGAGCAAAAACCGCTGGGATTGGGAACAGAAGAAAAAATGGATTCTGGAACTGAAACTGGGCAAACGCCCGATGGGCTGGAAAATGTGGGATTGGGAGCAGAAAAAGAAATGGATTAAAGGGAAACTGGATAAAGATCGCAACCTGGCGCGCAACCGCAAAAAAATTATGATGCGCTGGATTGAACGCCAAGAAGAAAAAATTAAAAAATGGGAACCGAGCAACCTGCCGAAAGGCATGAACGAAAAAACGAGCAAAAGCAAAAAACGCATGACCGCGAAAAAAATTGGCGTGCTGCGCAGCGTGAAACAGACCGTG

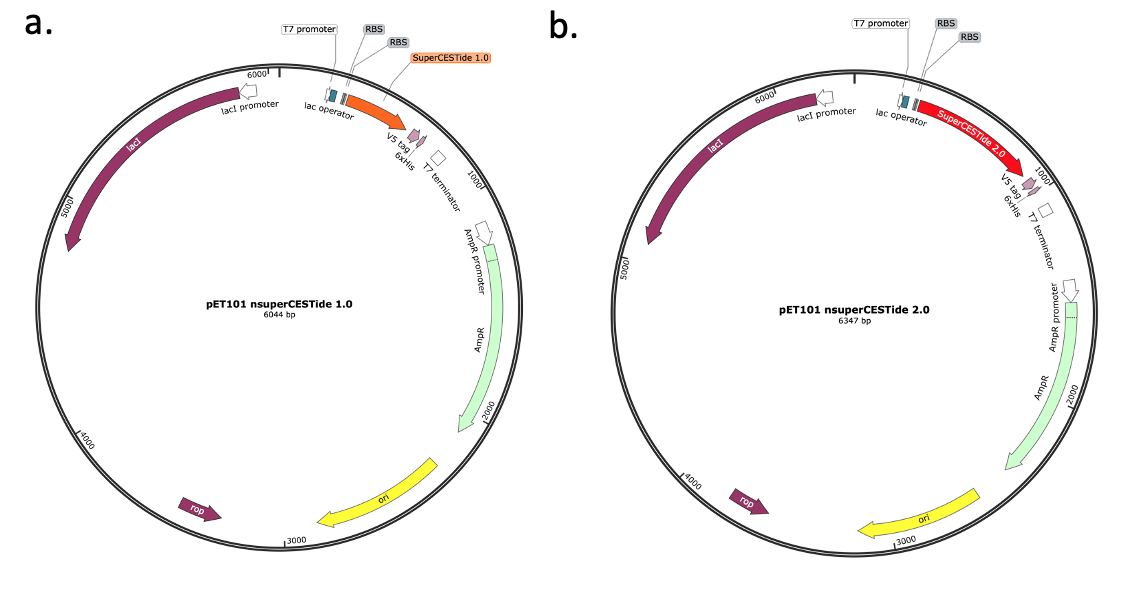

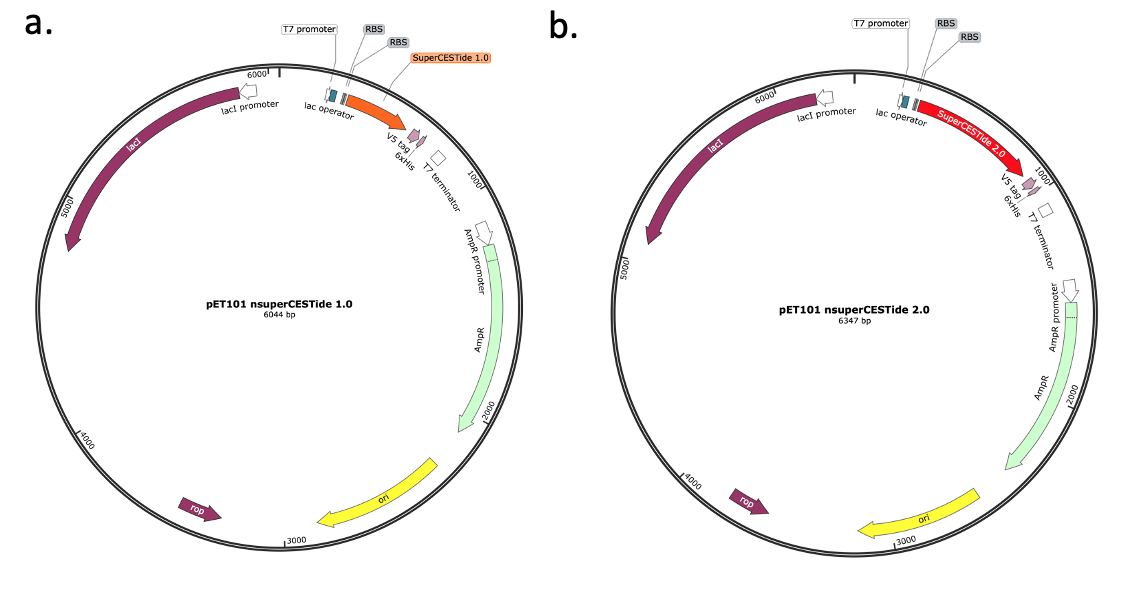

**Figure S1. Plasmid Maps of SuperCESTide 2.0 in the pET101 Vector. (a)** Plasmid map of superCESTide 2.0 in pET101, displaying the promoters, terminators, and synthetic gene in red, flanked by the ribosome binding sequences and affinity tags for various downstream processing.

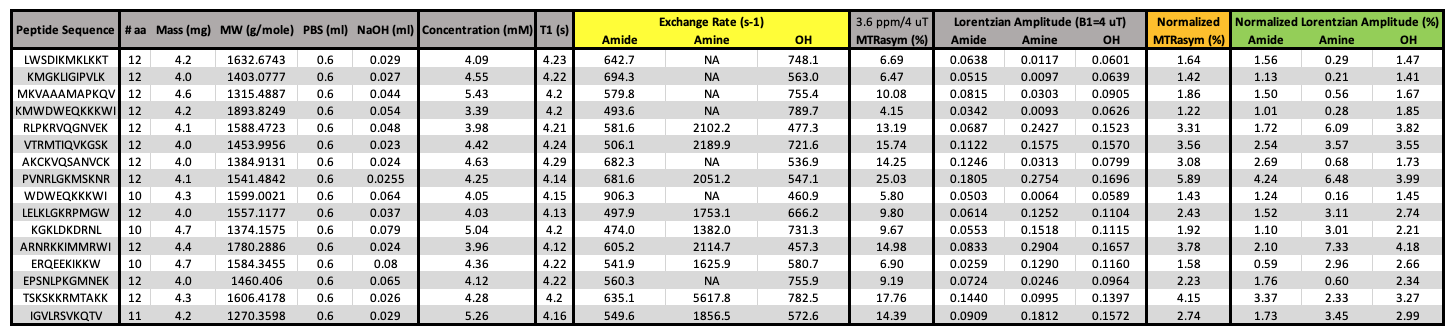
**Table S1. Exchange Rates and MTR_asym_ of Individual Peptides Used to Create the Synthetic Reporter Protein, SuperCESTide 2.0**

**6.2 – Purification using His-Tag affinity chromatography**

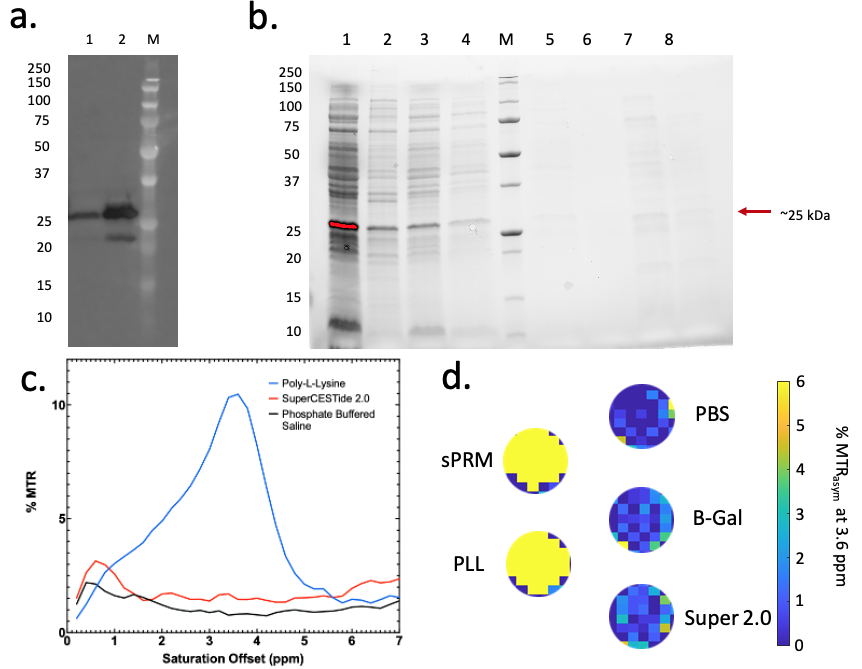
The following western blot and SDS-Page show successful protein expression at 22 °C, grown for 20 hours, and the following fractions from purification, respectively.

**Figure S2.** **Purification of superCESTide 2.0 with cobalt affinity chromatography resins.** **(a)** Western blot of soluble and insoluble fractions of superCESTide 2.0 grown in BL21 *E. coli*. Lane 1: Soluble superCESTide 2.0 grown at 22 deg C for 20 hrs, Lane 2: Insoluble superCESTide 2.0 grown at 22 deg C for 20 hrs, Lane M: Molecular Weight Marker. **(b)** SDS-PAGE fractions were obtained from the purification of superCESTide 2.0 at a higher concentration. Lane 1: Soluble fraction of 3x 25 mL cultures of superCESTide 2.0 grown at 22 deg C for 20 hrs, Lane 2: 1:10 dilution of the insoluble fraction of superCESTide 2.0 cultures grown at 22 deg C for 20 hrs, Lane 3: Flow through following pull down with cobalt resin, Lane 4, 5, 6: Wash steps involved in the purification of cobalt resin, Lanes 7, 8: Elution steps from purification. **(c)** MTR_asym_ of purified fractions of superCESTide 2.0 concentrated with 10 kDa amicon centrifugation filters. The red line represents purified protein at a concentration of 0.55 mg/mL; the blue line is of the control poly-L-lysine; the black line is phosphate-buffered saline as a control for the solvent. **(d)** ROIs of each well on phantom from MTR_aysm_ map generated by MRI, the max MTR_asym_ is 6% with yellow being the most intense signal.

**6.3 – Calibration of Size Exclusion Chromatography Column**

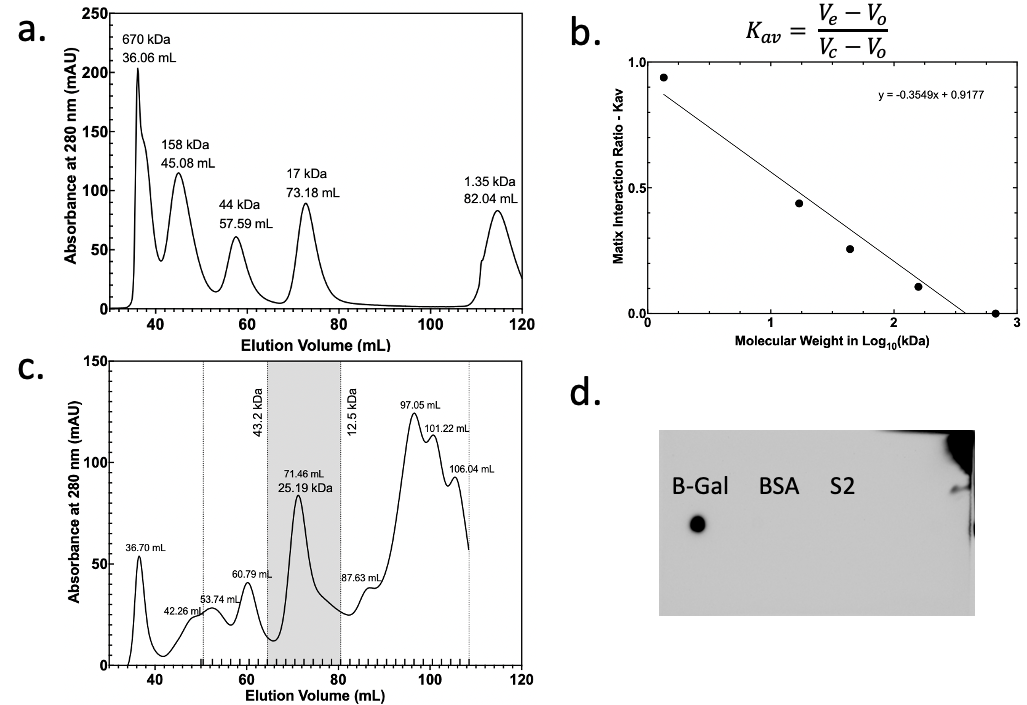

**Figure S3. Calibration of Size Exclusion Column and Initial Separation of SuperCESTide 2.0.** **(a)** Chromatogram collected of the Biorad gel filtration standards, at a flow rate of 0.5 mL/min, with a 100% PBS mobile phase. Each peak is denoted with the molecular weight of the protein from the standard and the volume at which the peak amount of protein is eluted. **(b)** The matrix interaction ratio (Kav) equation is displayed above the graph, where Ve is the elution volume of the protein of interest, Vo is the column's void volume determined by the retention volume of the largest protein, and V_c_ is the volume of the column. A linear regression was performed on K_av_ plotted against the log_10_ of the molecular weight, and the resulting linear function is shown on the graph. **(c)** Chromatogram collected from superCESTide 2.0 run on the SEC column using the same conditions used for calibration, so the linear equation could be used to calculate the approximate molecular weights of each peak. Each peak was denoted by its retention volume. The greyed peak was identified as having a molecular weight close to superCESTide 2.0, and the fractions were collected for this region spanning proteins with a size of 12.5 kDa to 43.2 kDa. **(d)** Dot blot of β-gal, BSA, and superCESTide 2.0 from SEC. β-gal was a positive control for 6-HIS tag binding, and BSA was a negative control for 6-HIS tag binding.
